# Hypoxia induces transcription of DOT1L in articular cartilage to protect against osteoarthritis

**DOI:** 10.1101/2021.05.14.444225

**Authors:** Astrid De Roover, Ana Escribano, Frederique M. F. Cornelis, Chahrazad Cherifi, Leire Casas-Fraile, An Sermon, Frederic Cailotto, Rik J. Lories, Silvia Monteagudo

**Author notes:** Correspondence to Professor Silvia Monteagudo, Laboratory of Tissue Homeostasis and Disease, Skeletal Biology and Engineering Research Center, Herestraat 49, O&N1 (10^th^ floor) – Box 813, 3000 Leuven, Belgium. RJL and SM are joint last authors.

## Abstract

Osteoarthritis is the most prevalent joint disease worldwide and a leading source of pain and disability. To date, this disease lacks curative treatment as underlying molecular mechanisms remain largely unknown. The histone methyltransferase DOT1L protects against osteoarthritis, and DOT1L-mediated H3K79 methylation is reduced in human and mouse osteoarthritic joints. Thus, restoring DOT1L function seems to be critical to preserve joint health. However, DOT1L-regulating molecules and networks remain elusive, in the joint and beyond. Here, we identify transcription factors and networks that regulate DOT1L gene expression using a novel bioinformatics pipeline. Thereby, we unravel an undiscovered link between the hypoxia pathway and DOT1L. We provide unprecedented evidence that hypoxia enhances DOT1L expression and H3K79 methylation via hypoxia-inducible factor-1 alpha (HIF-1α). Importantly, we demonstrate that DOT1L contributes to the protective effects of hypoxia in articular cartilage and osteoarthritis. Intra-articular treatment with a selective hypoxia mimetic in mice after surgical induction of osteoarthritis restores DOT1L function and stalls disease progression. Collectively, our data unravel a novel molecular mechanism that protects against osteoarthritis with hypoxia inducing DOT1L transcription in cartilage. Local treatment with a selective hypoxia mimetic in the joint restores DOT1L function and could be an attractive therapeutic strategy for osteoarthritis.

## INTRODUCTION

Osteoarthritis (OA) remains the most common chronic joint disease and a leading cause of disability with increasing incidence worldwide. It is characterised by progressive damage to the articular cartilage, varying degrees of synovial inflammation, subchondral bone remodelling, and osteophyte formation, leading to pain and loss of joint function.(1,2) Relevant molecular mechanisms with a role in the onset and progression of OA remain elusive. This may explain why current treatment is limited to symptom relief or joint replacement surgery and no disease-modifying therapy is available.

Restoring histone methyltransferase DOT1L may be an attractive strategy for therapy. The Disruptor of telomeric silencing 1-like (*DOT1L*) gene encodes an enzyme that methylates lysine 79 of histone H3 (H3K79), and is involved in epigenetic regulation of transcription.(3-5) Earlier, polymorphisms in *DOT1L* were associated with OA.(6,7) How DOT1L affects OA remained unknown until we identified DOT1L as master protector of cartilage health.(8) DOT1L activity, indicated by levels of methylated H3K79, is decreased in damaged areas from cartilage of OA patients compared to corresponding preserved areas and non-OA cartilage. Loss of DOT1L activity in human articular chondrocytes from healthy donors shifted their molecular signature towards an OA-like profile. In mice, intra-articular injection of a DOT1L inhibitor triggered OA. Heterozygous cartilage-specific *Dot1l* knockout (*Dot1l*^CART-KO^) mice spontaneously developed severe OA upon ageing.(9) Postnatal tamoxifen-induced conditional *Dot1l*^CART-KO^ mice developed more severe post-traumatic OA upon joint injury and spontaneous OA upon ageing compared to wild-type animals.(9) Mechanistically, DOT1L’s protective role is accomplished via restricting Wnt signalling, a pathway that when hyper-activated leads to OA and that is increasingly recognised as potential therapeutic target.(8,10,11)

Factors that regulate DOT1L levels and activity in the joint remain unknown. Targeting such mechanisms to maintain or restore DOT1L function appears to be a novel therapeutic opportunity to keep an optimal balance of Wnt signalling in cartilage, preserve joint health and inhibit progression of OA. Here, we aimed to discover DOT1L-regulating transcription factors and networks by analysing the human *DOT1L* promoter using an original bioinformatics pipeline. We identified a new mechanistic link between the hypoxia pathway and DOT1L, and validated this mechanism as therapeutic strategy to restore DOT1L function in articular cartilage and protect the joint against OA.

## RESULTS

### A novel bioinformatics pipeline identifies transcription factors regulating the *DOT1L* gene

To identify upstream signals regulating *DOT1L* gene expression, we conducted a bioinformatics analysis of the *DOT1L* promoter, using a novel pipeline we designed (Figure 1A). First, the DNA sequence of the human *DOT1L* proximal promoter was obtained from the eukaryotic promoter database (EPD).(12) We analysed this sequence with different bioinformatic tools, namely BindDB, PROMO, CONSITE and TFsitescan.(13-16). We found 276 transcription factors (TFs) predicted to interact with the *DOT1L* promoter *in silico*. As these bioinformatic tools use different algorithms and TF databases, we compared the outputs. This resulted in 31 TFs simultaneously predicted by at least 2 different tools, which were selected for further analysis (Figure 1A and B). To *in silico* interrogate binding of the 31 TFs to the *DOT1L* promoter, we analysed these individually using the Search Motif from the EPD website, as this tool predicts putative TF binding sites. This analysis corroborated putative binding of 29 TFs (Figure 1A).

**Figure 1.**
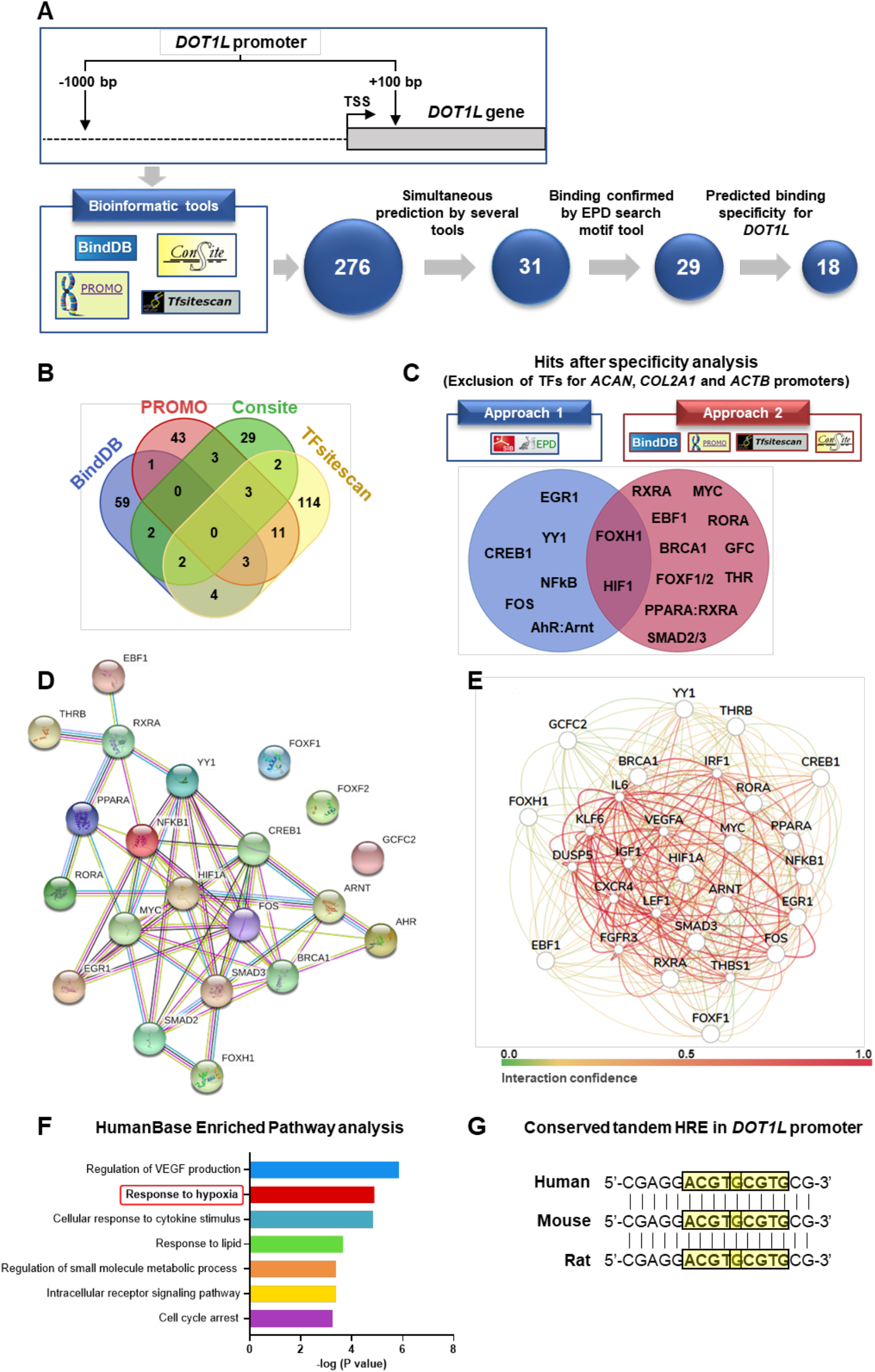
Bioinformatics analysis of the *DOT1L* promoter suggests an undiscovered link between hypoxia and DOT1L. (**A**) Overview of the bioinformatics analysis flow of the human *DOT1L* proximal promoter. The upper part of the panel displays the *DOT1L* gene promoter region that was used for the analysis, namely -1000 base pairs (bp) to +100 bp relative to the transcription start site (TSS). The lower part shows the four different bioinformatics web-based tools that were used and the transcription factor (TF) selection pipeline. (**B**) Venn diagram of the 276 TFs found by the four different tools, of which the TFs predicted by at least two different tools were selected for further analysis. (**C**) Overview of hits remaining after the specificity analysis. Two different approaches were used to determine whether the TFs were more specific for the *DOT1L* promoter compared to the *Aggrecan (ACAN), Collagen* (*COL2A1)* and *Actin* (*ACTB)* promoters. The diagram shows the 18 resulting TFs predicted to be more specific for *DOT1L* after exclusion of TFs by approach 1 or 2, and their overlap. (**D**) STRINGdb protein-protein network of the 18 resulting TFs upon the specificity analysis. (**E-F**) Cartilage-specific gene network of the 18 resulting TFs upon the specificity analysis using HumanBase (GIANT) (**E**) and its pathway enrichment analysis (**F**). (**G**) Presence of tandem hypoxia response elements (HREs) with consensus 5’-(A/G)CGTG-3’ (highlighted with yellow boxes) within the human, mouse and rat *DOT1L* gene promoters.

Then, we assessed potential specificity of these 29 TFs for *DOT1L*. We excluded TFs *in silico* predicted to bind to the promoters of genes that characterise the chondrocyte identity, namely *Aggrecan (ACAN)* and *Collagen2a1 (COL2A1)*, as well as housekeeping gene *Actin*, using two different approaches. The first used the EPD Search Motif tool to individually interrogate whether TFs predicted for *DOT1L* also bind to the promoter of the 3 mentioned control genes (Figure 1C). The second approach analysed the promoter sequences of the 3 control genes using the same bioinformatic tools as for *DOT1L*, and determined whether any of selected TFs appears in the output (Figure 1C). Combining both approaches resulted in 18 TFs that may selectively regulate *DOT1L* (Figure 1A and C). These TFs include Thyroid hormone receptor beta (THR), previously reported to increase *Dot1l* expression in tadpole intestine,(17) validating our novel in-house analysis pipeline.

### Bioinformatics analysis unravels an undiscovered link between hypoxia and *DOT1L*

Next, we explored interactions and regulatory networks of the obtained TFs. To this end, we used STRINGdb,(18) a database of known and predicted protein-protein interactions (Figure 1D). We also used HumanBase (GIANT) to build a cartilage-specific network (Figure 1E).(19) In the regulatory networks obtained, there was a prominent node around Hypoxia-inducible factor 1 alpha (HIF-1α) (Figure 1D and E) and the hypoxia pathway was enriched (Figure 1F). In our *in silico* specificity assessment (Figure 1C), HIF-1α was found to be specific for *DOT1L* by both approaches.

HIFs mediate the transcriptional response to low oxygen tension or hypoxia. They are heterodimeric TFs consisting of an unstable oxygen-sensitive α-subunit, and a stable β-subunit.(20) In the presence of oxygen, HIFα is degraded. Under hypoxia, HIFα is stabilised and dimerises with β-subunit. The heterodimers bind to hypoxia response elements (HREs) in the genome, regulating gene expression.

Adult articular cartilage is avascular and physiologically in a hypoxic state.(21) However, in OA, the hypoxic nature of cartilage is disrupted.(22,23) Mammals have three isoforms of α-subunit: HIF-1α, HIF-2α and HIF-3α.(24) Within human cartilage mainly HIF-1α and HIF-2α mediate transcriptional responses to hypoxia.(25) HIF-1α promotes cartilage homeostasis.(21,23,26) In contrast, HIF-2α is associated with chondrocyte hypertrophy and a catabolic response.(27,28)

HIFs can bind consensus sequence 5’-(A/G)CGTG-3’ within the HRE, but show differences in target gene specificity.(20,24,25,27) We identified HREs with consensus sequence 5’-(A/G)CGTG-3’ in the *DOT1L* promoter including overlapping tandem HREs (Figure 1G and Supplemental Figure 1). Two tandem HREs result in a stronger transcriptional response compared to one HRE.(29) Alignment of the human, mouse and rat *DOT1L* promoters revealed that the overlapping tandem HREs and surrounding region are conserved, highlighting the possible relevance of this regulatory motif (Supplemental Figure 2).

### Hypoxia increases *DOT1L* expression and H3K79 methylation in human articular chondrocytes

To investigate whether hypoxia regulates the *DOT1L* gene in articular cartilage, we studied effects of hypoxia mimetics or low oxygen levels on C28/I2 cells, a human articular chondrocyte cell line.(30) We used quantitative PCR to determine expression of *DOT1L* and positive control *Vascular endothelial growth factor* (*VEGF)*, a well-established hypoxia target gene. First, we treated C28/I2 cells with two pharmacological hypoxia mimetics. Treatment with IOX2 increased *DOT1L* expression in a concentration-dependent manner (R^2^=0.61, *p*=0.0002) (Figure 2A). Treatment with VH298 similarly led to increased *DOT1L* expression (R^2^=0.85, *p*<0.0001) (Figure 2B). Likewise, culturing C28/I2 cells in a hypoxia chamber promoted *DOT1L* expression [1.45-fold increase (95%CI: 1.18–1.80) *p*=0.0078] (Figure 2C). As expected, *VEGF* expression was increased in all experimental conditions [R^2^=0.87, *p*<0.0001; R^2^=0.97, *p*<0.0001; 3.12-fold increase (95%CI: 1.88 –5.18) *p*=0.0034; respectively] (Figure 2A-C).

**Figure 2.**
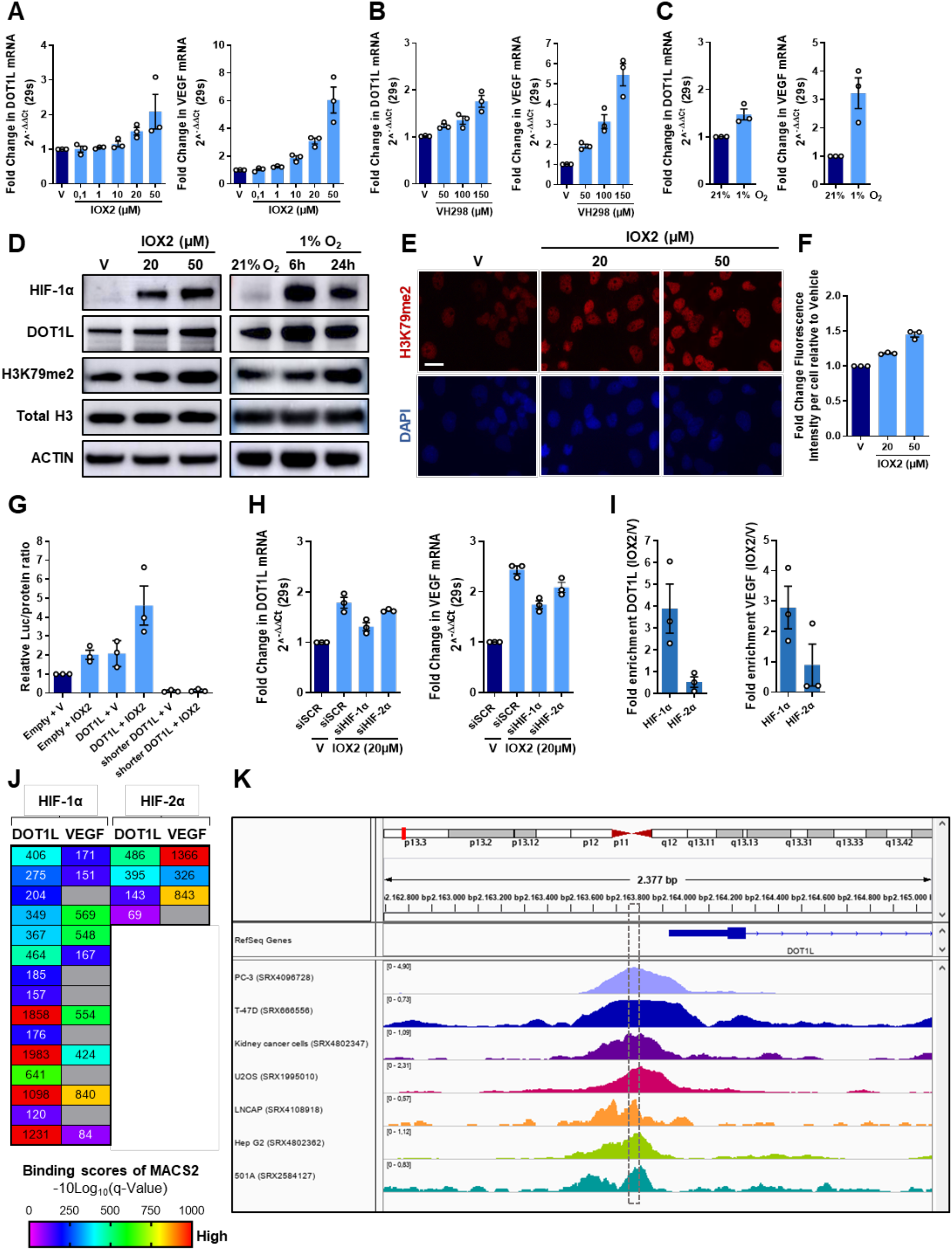
Hypoxia increases *DOT1L* expression and H3K79 methylation in human articular chondrocytes. (**A-B**) Real-time PCR analysis of *DOT1L* and *Vascular endothelial growth factor (VEGF)* after treatment with hypoxia mimetics IOX2 (**A**) and VH298 (**B**) or vehicle (V) at indicated concentrations for 72h in human articular chondrocyte C28/I2 cells [n=3 biologically independent experiments with 3 technical replicates, mean±sem., F_1,12_=28.32 *p*=0.0002, F_1,8_=56.91 *p*<0.0001 for *DOT1l*; F_1,12_=303.5 *p*<0.0001, F_1,8_=239.4 *p*<0.0001 for *VEGF* in (A) and (B) by Linear trend estimation ANOVA]. (**C**) Real-time PCR analysis of *DOT1L* and *VEGF* in C28/I2 cells upon culture in normoxic (21% O_2_) or hypoxic (1% O_2_) conditions for 6h (n=3, t_4_=4.94 *p*=0.0078 for *DOT1L*; t_4_=6.23 *p*=0.0034 for *VEGF* by t-test). (**D**) Immunoblot analysis of hypoxia-inducible factor 1 alpha (HIF-1α), DOT1L and H3K79 methylation amounts in C28/I2 cells treated with hypoxia mimetic IOX2 or V at the indicated concentrations for 96h and in response to hypoxia (1% O_2_) for the indicated times. The images are representative of images from two independent experiments. (**E-F**) Immunofluorescence staining analyses of H3K79 methylation (red) and DAPI (blue) in C28/I2 cells after treatment with IOX2 or V at the indicated concentrations for 72h. The images (**E**) are representative of 3 independent experiments with technical duplicates. Scale bar, 50 µm. Fluorescence intensity relative to V (**F**) was determined using 20 images per condition for each independent experiment (n=3, F_1,6_=300 *p*<0.0001 Linear trend estimation ANOVA). (**G**) Luciferase assay in C28/I2 cells transfected with either an empty plasmid, the full *DOT1L* promoter reporter plasmid or the shorter *DOT1L* promoter reporter plasmid, which does not contain the conserved overlapping tandem HREs, upon treatment with V or IOX2 (20µM) for 72h. Data are normalised to total protein concentration and relative to the empty plasmid and V condition (n=3, t_12_=2.625 *p*=0.0438 for DOT1L promotor+IOX2 vs. DOT1L promotor+V Sidak-corrected in one-way ANOVA with F_5,12_ = 55.97 *p*<0.0001). (**H**) Real-time PCR analysis of *DOT1L* and *VEGF* in C28/I2 cells after treatment with IOX2 (20µM) or V and siRNA-mediated silencing of *HIF-1α, HIF-2α* or scrambled control (siSCR) (n=3, t_8_= 9.530, 5.104, 1.427 – *p*<0.0001, =0.0028, =0.4715 for *DOT1L*; t_8_= 17.25, 6.568, 3.0426 – *p*<0.0001, =0.0051, =0.0473 for *VEGF*; V-siSCR vs IOX2-siSCR, IOX2-siSCR vs siHIF-1α, siHIF-1α vs siHIF-2α, Sidak-corrected in one-way ANOVA with F_3,8_=113 *p*<0.0001). (**I**) Chromatin-immunoprecipitation quantitative PCR (ChIP-qPCR) analysis of HIF-1α and HIF-2α binding to *DOT1L* and *VEGF* gene promoters in C28/I2 cells treated with hypoxia mimetic IOX2 (20 µM) for 72h versus V (n=3, t_2_=4.54 *p*=0.045, t_2_=1.51 *p*=0.271 for *DOT1L;* t_2_=3.76 *p*=0.064, t_2_=0.97 *p*=0.432 for *VEGF* by one sided t-test). (**J**) MACS2 binding scores of HIF-1α and HIF-2α around the *DOT1L* and *VEGF* transcription start site (TSS) of publicly available HIF-1α and HIF-2α ChIP-seqs on the ChIP-atlas database. (**K**) Visualisation of publicly available HIF-1α ChIP-seq data performed in various cell types around the *DOT1L* TSS. The dotted line box indicates the location of the overlapping HREs. All data were retrieved from ChIP-atlas and are mapped to the reference human genome (hg19) using the Integrative Genomics Viewer (IGV).

Then, we interrogated whether hypoxia also leads to increased DOT1L protein and H3K79 methylation, using western blot. Both IOX2 and incubation in a hypoxia chamber stabilised HIF-1α and increased DOT1L protein and H3K79 methylation in human articular chondrocytes (Figure 2D). Immunofluorescence further demonstrated increased DOT1L-mediated H3K79 methylation upon IOX2 treatment (R^2^=0.99, *p*<0.0001) (Figure 2E and F). Altogether, these data indicate that hypoxia enhances DOT1L expression and H3K79 methylation in articular chondrocytes.

### Hypoxia-mediated induction of *DOT1L* depends on HIF-1*α* but not HIF-2*α*

Next, we explored the molecular mechanism via which hypoxia induces *DOT1L* expression. First, we evaluated functionality of the conserved overlapping tandem HREs present in the *DOT1L* promoter, using a luciferase assay. To this end, the full human *DOT1L* promoter (−1000 bp to +91 bp relative from the TSS) and a shorter *DOT1L* promoter (−412 bp to +91 bp relative from TSS) in which the conserved overlapping tandem HREs were absent, were synthesised and cloned into the pGL3-basic vector upstream of a reporter luciferase gene (Supplemental Figure 3). These plasmids were transfected into C28/I2 cells followed by hypoxia mimetic IOX2 treatment. Luciferase activity was increased upon IOX2 stimulation in the full *DOT1L* promoter construct, but not the shorter promoter plasmid [2.21-fold (95%CI: 1.02-4.76) *p*=0.0438 in DOT1L full promoter condition] (Figure 2G). Thus, the conserved overlapping tandem HREs are functional and required for hypoxia-induced promoter activity.

Further, we interrogated the roles of HIF-1α and HIF-2α in *DOT1L* transcription. Silencing of *HIF-1α* blocked IOX2-mediated induction of *DOT1L* expression, while silencing *HIF-2α* had no effects in C28/I2 cells treated with hypoxia mimetic IOX2 [1.36-fold higher for siSCR vs siHIF-1α (95%CI: 1.13-1.63) *p*=0.0028 for *DOT1L*, 1.4-fold higher for *VEGF* (95%CI: 1.20-1.63) *p*=0.0005] (Figure 2H and Supplemental Figure 4). Chromatin immunoprecipitation– qPCR (ChIP-qPCR) in C28/I2 cells treated with IOX2 demonstrated that HIF-1α localised at the *DOT1L* gene promoter, but not HIF-2α [3.59-fold (95%CI: 1.07-12.09) *p*=0.045 for *HIF-1A*] (Figure 2I). To further confirm this molecular mechanism, we interrogated *DOT1L* as a potential target gene of HIF-1α and HIF-2α using ChIP-Atlas,(31) an integrative and comprehensive data-mining suite of public ChIP-seq data. This analysis revealed relatively higher MACS2 scores for HIF-1α compared to HIF-2α indicating higher binding to *DOT1L* (Figure 2J). Binding of HIF-1α to the *DOT1L* promoter was found around the area of the overlapping HREs in multiple ChIP-atlas datasets (Figure 2K). Collectively, these data demonstrate that hypoxia directly regulates *DOT1L* expression via HIF-1α.

### DOT1L contributes to the protective effects of hypoxia in human articular chondrocytes

To translationally validate these findings, we assessed effects of hypoxia on primary human articular chondrocytes (hACs). Also in these cells, IOX2 treatment and culture in a hypoxia chamber induced *DOT1L* expression [R^2^=0.49, *p*=0.006; 1.80-fold increase (95%CI: 1.10– 2.94) *p* = 0.0296; respectively] (Figure 3A and B). Expression of positive control VEGF was also induced [R^2^=0.91, *p*<0.0001; 7.27-fold increase (95%CI: 4.18–12.63) *p*=0.0006; respectively] (Figure 3A and B). Then, we verified that a hypoxic environment is beneficial for the molecular phenotype of the articular chondrocyte. Hypoxia mimetic IOX2 increased expression of *COL2A1* and *ACAN* (R^2^=0.97, *p*=0.0004; R^2^=0.72, *p*<0.0001; respectively) (Figure 3C). Also, culturing primary hACs in a hypoxia chamber induced healthy chondrocyte markers [11.36-fold increase for *COL2A1* (95%CI: 7.12–18.12) *p*=0.0001; 4.14-fold for *ACAN* (95%CI: 3.23–5.31) *p*<0.0001] (Figure 3D).

**Figure 3.**
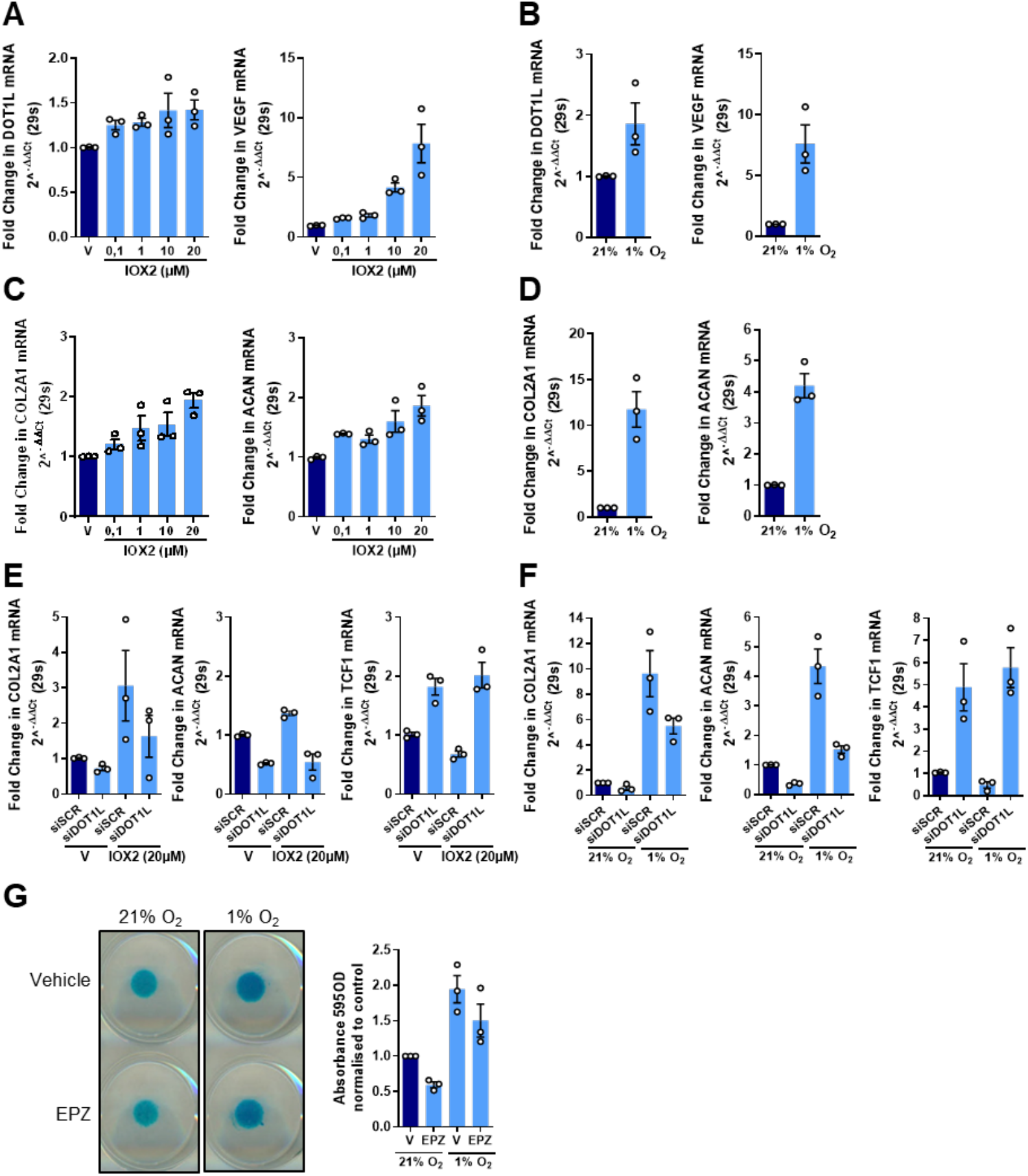
DOT1L contributes to the protective effects of hypoxia in human articular chondrocytes. (**A**) Real-time PCR analysis of *DOT1L* and *VEGF* in primary human articular chondrocytes (hACs) after treatment with hypoxia mimetic IOX2 or vehicle (V) at the indicated concentrations for 72h (n=3, F_1,10_=12.05 *p*=0.006 for *DOT1l;* F_1,10_=213.9 *p*<0.0001 for *VEGF* by Linear trend estimation ANOVA). (**B**) Real-time PCR analysis of *DOT1L* and *VEGF* in primary hACs cultured in normoxic (21% O_2_) or hypoxic (1% O_2_) conditions for 14 days. (n=3, t_4_=3.312 *p*=0.0296 for *DOT1L*; t_4_=9.966 *p*=0.0006 for *VEGF* by t-test). (**C**) Real-time PCR analysis of *COL2A1* and *ACAN* in primary hACs after treatment with IOX2 at the indicated concentrations or V for 72h (n=3, F_1,10_=27.85 *p*=0.0004 for *COL2A1*; F_1,10_=41.02 *p*<0.0001 for *ACAN* by Linear trend estimation ANOVA). (**D**) Real-time PCR analysis of *COL2A1* and *ACAN* in primary hACs cultured in normoxic (21% O_2_) or hypoxic (1% O_2_) conditions for 14 days (n=3, t_4_=14.45 *p*=0.0001 for *COL2A1*; t_4_=15.86 *p*<0.0001 for *ACAN* by t-test). (**E**) Real-time PCR analysis of *COL2A1, ACAN* and *TCF1* in primary hACs after treatment with hypoxia mimetic IOX2 (20 µM) or V and siRNA-mediated silencing of *DOT1L* (siDOT1L) or scrambled control (siSCR) for 72h (n=3, t_8_=0.7729 *p*=0.7103, t_8_=1.658 *p*=0.2532 siSCR vs siDOT1L with V and IOX2 for *COL2A1* Sidak-corrected in two-way ANOVA with F_1,8_=6.332 *p*=0.0360 for treatment status, and F_1,8_=2.956 *p*=0.1239 for silencing status; t_8_=3.001 *p*=0.0338, t_8_=4.623 *p*=0.0034 siSCR vs siDOT1L with V and IOX2 for *ACAN* Sidak-corrected in two-way ANOVA with F_1,8_=0.8014 *p*=0.3968 for treatment status, and F_1,8_=123.4 *p*=0.0007 for silencing status; t_8_=5.314 *p*=0.0014, t_8_=9.885 *p*<0.0001 siSCR vs siDOT1L with V and IOX2 for *TCF1* Sidak-corrected in two-way ANOVA with F_1,8_=10.45 *p*=0.0120 for interaction between treatment and silencing status). (**F**) Real-time PCR analysis of *COL2A1, ACAN* and *TCF1* in primary hACs cultured in normoxic (21% O_2_) or hypoxic (1% O_2_) conditions for 14 days and siRNA-mediated silencing of *DOT1L* or scrambled control (siSCR) (n=3, t_8_=2.453 *p*=0.2159, t_8_=2.194 *p*=0.3082 siSCR vs siDOT1L in normoxia and hypoxia for *COL2A1* Sidak-corrected in two-way ANOVA with F_1,8_=168.8 *p*<0.0001 for oxygen status, and F_1,8_=10.80 *p*=0.011 for silencing status; t_8_=7.799 *p*=0.0003, t_8_=7.909 *p*=0.0003 siSCR vs siDOT1L in normoxia and hypoxia for *ACAN* Sidak-corrected in two-way ANOVA with F_1,8_=240.5 *p*<0.0001 for oxygen status, and F_1,8_=123.4 *p*<0.0001 for silencing status; t_8_=4.713 *p*=0.0091, t_8_=8.119 *p*=0.0002 siSCR vs siDOT1L in normoxia and hypoxia for *TCF1* Sidak-corrected in two-way ANOVA with F_1,8_=5.803 *p*=0.0426 for interaction between oxygen and silencing status). (**G**) Alcian blue staining of primary hACs micromasses cultured in normoxic (21% O_2_) or hypoxic (1% O_2_) conditions treated with V or DOT1L inhibitor EPZ-5676 (EPZ) for 14 days. The images are representative of 3 independent experiments with technical triplicates. Quantification of staining relative to V in normoxic conditions was determined by colorimetry at 595nm (n=3, t_8_=3.868 *p*=0.0282, t_8_=1.992 *p*=0.382 EPZ vs V in normoxia or hypoxia, t_8_=4.812 *p*=0.008, t_8_=6.688 *p*=0.0009 normoxia-V vs hypoxia-V or EPZ Sidak-corrected in two-way ANOVA with F_1,8_=66.13 *p*<0.0001 for oxygen status, F_1,8_=17.17 *p*=0.0032 for treatment status, and F_1,8_=1.759 *p*=0.2213 for interaction between oxygen and treatment status).

Our data demonstrate that hypoxia induces DOT1L expression and has protective effects in primary hACs. A key question is whether DOT1L contributes to hypoxia’s protective effects. To answer this, we evaluated effects of IOX2-induced hypoxia in the absence of DOT1L in primary hACs. Silencing of *DOT1L* using siRNA partially blocked protective effects of hypoxia on *ACAN* expression [1.72-fold (95%CI: 0.83-3.55) *p*=0.1239 for *COL2A1*; 2.29-fold (95%CI: 1.61-3.26) *p*=0.0007 for *ACAN*] (Figure 3E). We earlier demonstrated that DOT1L’s protective role is exerted via limiting Wnt signalling.(8,9) We assessed Wnt signalling activity in IOX2-treated hACs by measuring expression of *TCF1*, a direct Wnt target gene epigenetically regulated by DOT1L.(8) IOX2 treatment reduced Wnt signalling activity, and *DOT1L* silencing impaired this effect (effect of silencing and oxygen exposure showed interaction *p*=0.0120) (Figure 3E and Supplemental Figure 5). This indicates that DOT1L participates in effects of hypoxia on Wnt signalling reduction. Silencing of DOT1L in primary hACs cultured in a hypoxia chamber reduced expression of healthy chondrocyte markers and increased Wnt target gene *TCF1* [1.77-fold (95%CI: 1.19-2.64) *p*=0.011 for *COL2A1*; 2.79-fold (95%CI: 2.25-3.46) *p<*0.0001 for *ACAN*; for *TCF1* effect of silencing and oxygen exposure showed interaction *p*=0.0426] (Figure 3F and Supplementary Figure 5). To further confirm this, we used three-dimensional micromass cultures of primary hACs. Culturing these micromasses under hypoxia increased glycosaminoglycan content as determined by Alcian blue staining [1.71-fold (95%CI: 1.47-2.01) p<0.001], while DOT1L inhibition with EPZ-5676 (EPZ) partially blocked this increase [0.75-fold (95%CI: 0.65-0.88) p=0.032] (Figure 3G). Collectively, these results demonstrate that DOT1L contributes to the protective effects of hypoxia in articular cartilage.

### Intra-articular treatment with IOX2 halts OA in mice and restores DOT1L in articular cartilage

We investigated therapeutic implications of our findings for OA. We previously showed that DOT1L and H3K79 methylation are reduced in human and mouse OA cartilage compared to non-OA cartilage.(8,9) Also, HIF-1α is reduced in OA cartilage.(22,23,27) Yet, to our knowledge, local administration of selective hypoxia mimetics has not been evaluated in a clinically relevant mouse model of OA. We investigated effects of a hypoxia mimetic on OA using the destabilisation of the medial meniscus (DMM) mouse disease model.(32,33) Before initiating *in vivo* treatments, we corroborated that DMM-operated mice showed a concomitant down-regulation in DOT1L and HIF-1α proteins in articular cartilage compared to sham-operated controls (Supplemental Figure 6). Upon these observations we proceeded with the *in vivo* pharmacological intervention. Starting one week after DMM surgery, mice were intra-articularly injected with IOX2 every 10 days, and knee joints were collected 12 weeks after surgery (Figure 4A). Histological analysis showed that IOX2 treatment reduced cartilage damage [difference of means between IOX2-treated and vehicle in DMM 1.13 (95%CI: 0.19-2.08) *p*=0.0202], osteophyte formation [difference of means 0.40 (95%CI: 0.07-0.72) *p*=0.0172] and synovial inflammation [difference of means 0.36 (95%CI: 0.001-0.72) *p=*0.049] upon DMM (Figure 4B-D). Immunohistochemistry showed that IOX2 treatment effectively rescued HIF-1α, and increased DOT1L expression and H3K79 methylation after DMM (Figure 4E-G). These *in vivo* data indicate that intra-articular treatment with a selective hypoxia mimetic restores DOT1L and H3K79 methylation, and protects against OA.

**Figure 4.**
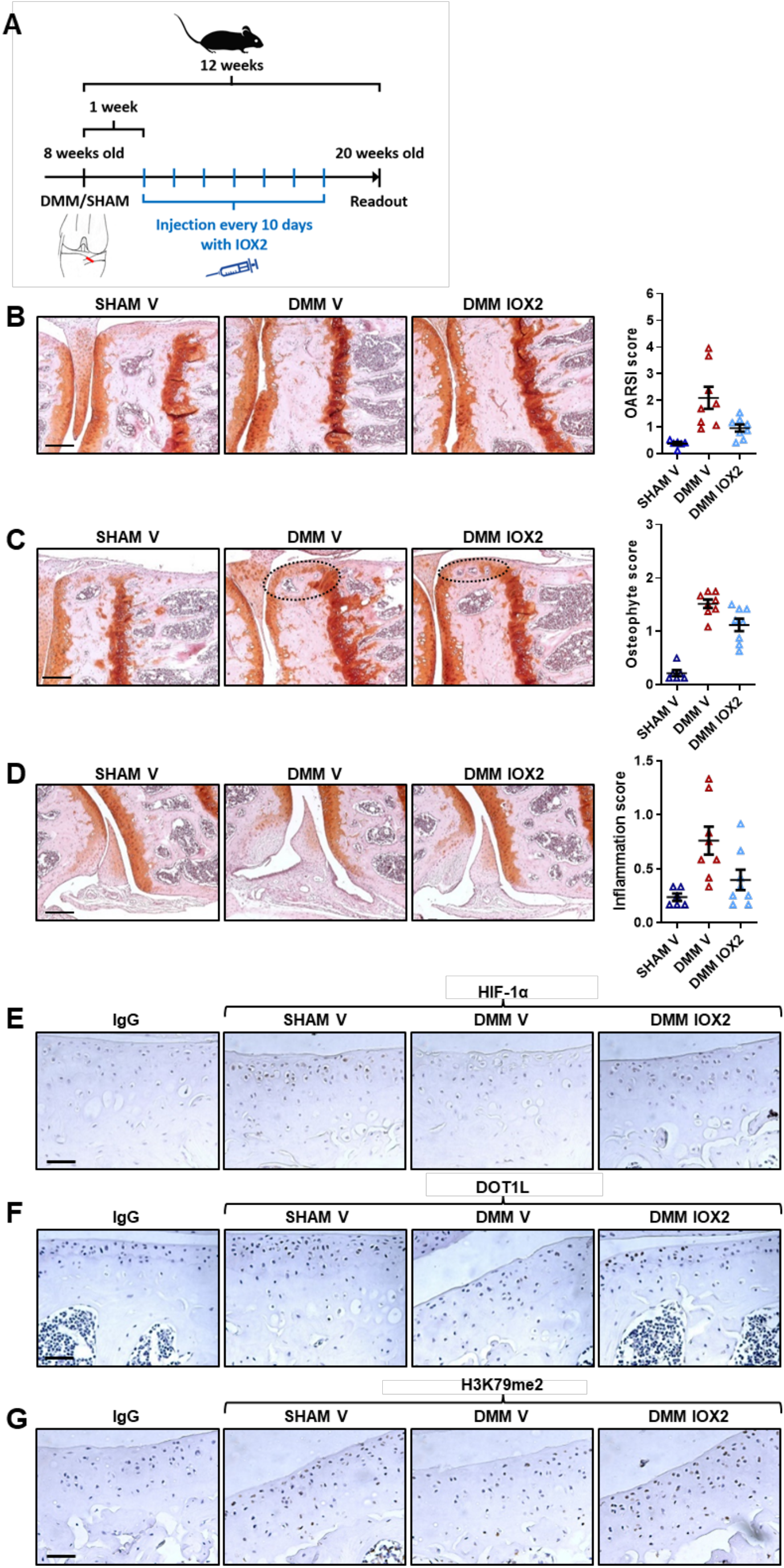
Intra-articular injection with IOX2 halts OA in mice and restores DOT1L in articular cartilage. (**A**) Time course of intra-articular injections with IOX2 or vehicle (V) in DMM or Sham operated wild-type mice. (**B**) Frontal hematoxylin-safraninO staining of the medial tibia and femur and quantification of articular cartilage damage at the four quadrants, evaluated by OARSI score. Scale bar, 200 µm. Data are presented as individual data points (mean) [n=6 (SHAM V) and n=8 (DMM V and DMM IOX2), t_19_=4.257 *p*=0.0013, t_19_=3.037 *p*=0.0202 SHAM V vs DMM V and DMM V vs DMM IOX2, Sidak-corrected in one-way ANOVA with F_2,19_=9.784 *p*=0.0012]. (**C**) Frontal hematoxylin-safraninO staining of the medial tibia and femur and quantification of osteophytes at the medial tibia and femur. Scale bar, 200 µm. Data are presented as individual data points [n=6 (SHAM V) and n=8 (DMM V and DMM IOX2), t_19_=9.456 *p*<0.0001, t_19_=3.109 *p*=0.0172 SHAM V vs DMM V and DMM V vs DMM IOX2, Sidak-corrected in one-way ANOVA with F_2,19_=45.73 *p*<0.0001]. (**D**) Frontal hematoxylin-safraninO staining of the lateral synovium and quantification of inflammation. Scale bar, 200 µm. Data are presented as individual data points (mean) [n=6 (SHAM V) and n=8 (DMM V and DMM IOX2), t_19_=3.494 *p*=0.0073, t_19_=2.624 *p*=0.0497 SHAM V vs DMM V and DMM V vs DMM IOX2, Sidak-corrected in one-way ANOVA with F_2,19_=6.750 *p*=0.0061]. (**E-F-G**) Immunohistochemical detection of HIF-1α, DOT1L and H3K79me2 in the articular cartilage of wild-type mice treated with IOX2 or V upon OA triggered by DMM surgery compared to sham operated mice. The images are representative of three different animals. Scale bar, 50 µm.

## DISCUSSION

This study evidences that restoring hypoxia in the joint could be an attractive therapeutic strategy for OA, since it rescues DOT1L activity in cartilage. Despite DOT1L’s fundamental role in diverse biological processes, to date little is known about how DOT1L expression and activity are regulated.(34,35) We identified potential TFs that regulate the *DOT1L* gene using a new bioinformatics analysis. Several applications developed to predict regulatory elements have poor predictive specificity.(36) Our novel pipeline that includes and compares multiple available online tools could effectively identify regulators of the *DOT1L* gene. Hence, our pipeline design may be used to identify regulators for any gene of interest.

The avascular cartilage is in a permanent hypoxic state throughout life. Evolutionarily, articular chondrocytes are well adapted to hypoxia and low oxygen levels maintain cartilage homeostasis.(37-39) Our data indicate that hypoxia is disrupted in OA, in agreement with recent studies demonstrating increased oxygen concentrations in OA cartilage.(22,23) OA-associated cartilage damage may allow deeper synovial fluid penetration and, as a consequence, more oxygen supply to the cartilage. Another hypothesis suggests that OA synovial membrane inflammation and hyper-vascularity may alter the oxygen diffusion characteristics resulting in higher oxygen concentration in synovial fluid and thus in cartilage.

Within human cartilage, mainly HIF-1α and HIF-2α mediate the response to hypoxia. HIF-1α has been reported to promote cartilage homeostasis in several ways. For instance, HIF-1α increases expression of anabolic chondrogenic genes, such as *COL2A1* and *ACAN*.(37) In addition, HIF-1α supresses catabolic proteins, such as *Matrix metallopeptidase 13 (MMP-13)* and *Nuclear factor-kappa B* (*NFkB*).(22,37) In contrast, the role of HIF-2α is still controversial and mainly associated with chondrocyte hypertrophy and a catabolic response.(27,28) Our present data demonstrating DOT1L is regulated by HIF-1α and not by HIF-2α are in line with HIF-1α’s established protective role.

Hypoxia promotes chondrogenic matrix genes while supressing catabolic enzymes,(39) however, underlying mechanisms remain incompletely defined. Duval *et al*. demonstrated that hypoxia induces chondrogenesis in mesenchymal stem cells by HIF-1α binding to the *SOX9* promoter, subsequently increasing *COL2A1* expression.(40) Bouaziz *et al*. reported that HIF-1α can interact with β-catenin thereby reducing the binding of TCF4 to Wnt target gene promoters.(23) Thus, we currently lack a complete understanding of oxygen sensitive pathways and the response to hypoxia, in particular in articular cartilage. We reveal that DOT1L contributes to the protective effects of hypoxia. Silencing or inhibiting *DOT1L* reduced beneficial effects of hypoxia on primary hACs in monolayer and three-dimensional cultures. Considering DOT1L’s role in Wnt signalling regulation, we assessed effects on Wnt activity, demonstrating that hypoxia limits Wnt signalling. This is in line with findings of Bouaziz *et al*.(23) However, we also demonstrate that this effect on Wnt is impaired upon *DOT1L* silencing.

We identified a regulatory motif in the *DOT1L* promoter consisting of two overlapping HRE tandem repeats, conserved among species. Due to steric hindrance, it is unlikely that both HREs are functional. Yet, Fukasawa *et al*. previously demonstrated that two tandem HREs result in a stronger transcriptional response compared to only one.(29) Nevertheless, the presence of a motif does not necessarily mean that the TF will bind. Our luciferase assay results demonstrate the functionality of the overlapping tandem HREs present in the *DOT1L* promoter.

As mentioned, hypoxia increases healthy chondrocyte markers and reduces catabolic markers in normal and OA chondrocytes *in vitro*.(39) Yet, to date, there are no reports about successful *in vivo* pharmacological interventions to locally restore hypoxia in OA joints. To our knowledge, only two studies investigated whether stabilisation of HIF-1α could prevent OA.(41,42) Both studies used dimethyloxalylglycine (DMOG), a 2-oxoglutarate (2-OG) analogue that acts as a broad spectrum inhibitor against all 2-OG-dependent dioxygenases. Notably, 2-OG-dependent dioxygenases have multiple roles in cell biology, participating in oxygen sensing, lipid metabolism, collagen and carnitine biosynthesis and histone demethylation.(43-47) Gelse *et al*. injected DMOG intra-articularly in knees of 8-week-old STR/ort mice.(41) This treatment did not ameliorate spontaneous OA, which was explained by the lack of specificity of DMOG, which interferes with collagen biosynthesis resulting in reduced *COL2A1* expression, and induces catabolic cytokines.(41) Hu *et al*. performed intraperitoneal injections of DMOG in 10-week-old DMM mice every day.(42) Although this approach seemed to reduce cartilage damage, an important point of attention that would limit its clinical application is that such systemic approach using a broad spectrum inhibitor (DMOG) would trigger a systemic inhibition of 2-OG-dependent dioxygenases. Therefore, this setup could be regarded as not hypoxia nor tissue selective. Here, intra-articular administration of IOX2, a selective hypoxia mimetic, was able to halt OA in a relevant mouse OA model.

Importantly, the discovery that hypoxia controls the transcription of the *DOT1L* gene might have implications in diseases beyond OA. Of note, our ChIP-atlas analysis showed that HIF-1α binds to the *DOT1L* promoter in several cell types. This might indicate that the regulatory mechanism identified here could indeed play a role in organs and tissues beyond cartilage. Based on literature, hypoxia and DOT1L are important in several coinciding processes throughout the human body, thereby suggesting that a hypoxia-mediated regulation of DOT1L could be involved. For example, both hypoxia and DOT1L have been reported to be essential during early erythropoiesis(48,49). In addition, hypoxia and DOT1L are both linked to the immune response and to kidney injury(50-55). Also in cancer, hypoxia and DOT1L play important roles(51,56-58). In conclusion, our study identifies that hypoxia regulates the expression of *DOT1L* via HIF-1α. We demonstrate that DOT1L contributes to the protective effects of hypoxia on OA. Translationally, local treatment with a selective hypoxia mimetic halts disease progression in the DMM mouse OA model, demonstrating that targeting hypoxia could be an attractive therapeutic strategy for OA.

## METHODS

### Materials

The hypoxia mimetics IOX2 and VH298 were purchased from Sigma and Tocris, respectively.

### Bioinformatics analysis

The *DOT1L* proximal promoter sequence (1000 bp upstream and 100 bp downstream relative to the transcription start site) was obtained from the online tool Eukaryotic Promoter Database (EPD) (https://epd.epfl.ch//index.php).(12) This DNA sequence was analysed using online freely available web-based tools to predict transcription factors (TFs), namely CONSITE (http://consite.genereg.net),(15) TFsitescan (http://www.ifti.org/cgi-bin/ifti/Tfsitescan.pl),(16) BindDB (http://bind-db.huji.ac.il),(13) and PROMO (http://alggen.lsi.upc.es/cgi-bin/promo_v3/promo/promoinit.cgi?dirDB=TF_8.3).(14) We compared the outputs from the different online tools. Only TFs that were predicted by at least 2 different tools were selected for further analysis. Next, we used the EPD Search Motif tool, which uses the JASPAR database, to confirm predicted binding to the *DOT1L* proximal promoter. Potential specificity for *DOT1L* was *in silico* interrogated by assessing the binding of these TFs to the promoters of *Aggrecan, Collagen2a1* and *Actin* using two different approaches. The first approach consisted of using the EPD Search Motif tool to individually interrogate whether the selected TFs predicted to bind to the *DOT1L* gene, also bind to the promoter of the control genes mentioned above. The second approach consisted of analysing the promoter sequences of the control genes using CONSITE, TFsitescan, BindDB and PROMO, and determine whether any of the selected TFs appears in the output. Finally, STRINGdb (https://string-db.org)(18) and HumanBase (https://hb.flatironinstitute.org)(19) were used to explore protein-protein interactions and cartilage specific regulatory networks of the predicted TFs, respectively.

### Cell culture

Human immortalised articular chondrocyte C28/I2 cells were purchased from Merck Millipore and cultured in DMEM/F12 (Gibco) containing 10% fetal bovine serum (FBS) (Gibco), 1% (vol/vol) antibiotic/antimycotic (Gibco) and 1% L-glutamine (Gibco) in a humidified atmosphere at 37 °C and 5 % CO_2_. Primary human primary articular chondrocytes (hACs) were isolated from patients undergoing hip replacement surgery for osteoporotic or malignancy-associated fractures with informed consent and ethical approval by the University Hospital Leuven Ethics Committee. First, cartilage was dissected from the joint surface, rinsed with PBS and cut into small pieces. The cartilage pieces were incubated with 2 mg/ml pronase solution (Roche) for 90 min at 37 °C and digested overnight at 37 °C in 1.5 mg/ml collagenase B solution (Roche). Then, the preparation was filtered through a 70 µM strainer and cells were plated in culture flasks and cultured in a humidified atmosphere at 37 °C and 5% CO_2_. Culture medium consisted of DMEM/F12 (Gibco), 10% FBS (Gibco), 1% (vol/vol) antibiotic/antimycotic (Gibco) and 1% L-glutamine (Gibco).

### Small Interfering RNA Transfection

Cells were transfected with lipofectamin RNAiMAX (Invitrogen) as transfection reagent, together with non-targeting siGENOME siRNA (siSCR) or siGENOME siRNA against DOT1L, HIF-1α or HIF-2α (Dharmacon) following the protocols provided by the manufacturer.

### Quantitative PCR

Total RNA was extracted using the Nucleospin RNA II kit (Macherey-Nagel). cDNA was synthesised using the RevertAidHminus First Strand cDNA synthesis kit (Thermo Fisher Scientific) according to the manufacturers’ guidelines. Quantitative PCR analyses were carried out as described previously using Maxima SYBRgreen qPCR master mix system (Thermo Fisher Scientific).(8) Gene expression was calculated following normalisation to housekeeping gene S29 mRNA levels using the comparative Ct (cycle threshold) method. The following PCR conditions were used: incubation for 10 min at 95 °C followed by 40 amplification cycles of 15 s of denaturation at 95 °C followed by 45 s of annealing-elongation at 60 °C. Melting curve analysis was performed to determine the specificity of the PCR. Primers used for qPCR analysis are listed in Supplemental Table 1.

### Cell lysis and western blotting

Cells were lysed in IP Lysis/Wash buffer (Thermo Fisher) supplemented with 5% (vol/vol) Protease Mixture Inhibitor (Sigma) and 1 mM phenylmethanesulfonyl (Sigma). After two homogenisation cycles (7 s) with an ultrasonic cell disruptor (Microson; Misonix), total cell lysates were centrifuged at 18,000 g for 10 min. The supernatant containing proteins was collected and the protein concentration was determined by Pierce BCA Protein Assay Kit (Thermo Scientific). Immunoblotting analysis was carried out as previously described.(8) Antibodies against Actin (Sigma, A2066; dilution 1:4,000), DOT1L (Cell Signaling, #77087; dilution 1:1,000), HIF-1α (Abcam, ab82832; dilution 1:1,000), total H3 (Abcam, ab1791; dilution 1:10,000) and H3K79me2 (Abcam, ab3594; dilution 1:1,000) were used following manufacturer’s instructions. The blotting signals were detected using the SuperSignalWest Femto Maximum Sensitivity Substrate system (Thermo Scientific).

### ChIP analysis

Chromatin immunoprecipitation (ChIP) assays were performed using the Agarose ChIP kit (Thermo Fisher Scientific) according to the manufacturer’s recommendations. Briefly, cell samples were cross-linked with 1% formaldehyde for 10 min. This reaction was stopped by adding glycine to a 125 mM final concentration. The fixed cells were lysed and the chromatin was fragmented by nuclease digestion. Further, the sheared chromatin was incubated with antibodies against HIF-1α (Abcam, ab1; dilution 1:50) and HIF-2α (Abcam, ab199; dilution 1:50) and recovered by binding to protein A/G agarose. Eluted DNA fragments were used directly for qPCR. Primers used for ChIP-qPCR analysis are listed in Supplemental Table 2. Bioinformatics *in silico* ChIP analysis was performed using the ChIP-Atlas (https://chip-atlas.org)(31), an integrative and comprehensive data-mining suite of public ChIP-seq data. The feature Target Genes was used to predict target genes bound by the given TFs HIF-1α and HIF-2α. From these results, the individual ChIP-seq experiments that showed binding to *DOT1L* were selected for further analysis. The peak-caller Model-based Analysis of ChIP-seq (MACS) algorithm captures the influence of genome complexity to evaluate the significance of enriched ChIP regions. These MACS2 binding significance scores were evaluated for *DOT1L* and *VEGF* in each individual ChIP-seq experiment. Finally, BigWig data of HIF-1α ChIP-seq perfomed in several cell types were visualised around the *DOT1L* transcription start site. All the data were mapped to the reference human genome (hg19) using the Integrative Genomics Viewer (IGV).

### Luciferase reporter assay

The full *DOT1L* promoter sequence (−1000 bp to +91 bp relative to TSS) and a shorter promoter sequence (−412 bp to +91 bp relative to TSS) in which the conserved overlapping tandem HREs were removed, were amplified by PCR and cloned into the pGL3-Basic luciferase reporter vector (Promega, E1751) using the KpnI and XhoI sites [promoter sequence defined using the Eukaryotic promoter database (https://epd.epfl.ch//index.php)]. Primers used for the amplification are described in Supplemental Table 3. C28/I2 cells were seeded in 24-well plates. After 24h, the cells were transfected with the full or shorter promoter reporter plasmids using Lipofectamine LTX Reagent with PLUS Reagent (Invitrogen) according to the manufacturer’s protocol. After 24h, the cells were stimulated with vehicle (DMSO) or IOX2 (20 µM) for 72h. The luciferase activity was assessed with Luciferase Assay System (Promega). As a control, the total protein concentration was determined by the Pierce BCA Protein Assay Kit (Thermo Scientific). Finally, the ratio of Firefly luciferase to total protein was determined as relative luciferase activity.

### Immunofluorescence

C28/I2 cells were seeded in Nunc(tm) Lab-Tek(tm) II (ThermoFisher) chamber slides. The following day, the cells were treated with 20 µM IOX2, 50 µM IOX2 or vehicle DMSO for 72h. Then, the cells were fixed using 3.7% Formaldehyde in PBS for 10 min and antigen retrieval was performed using 1% SDS in PBS for 2 min. The cells were blocked in 1% BSA for 30 min and incubated with primary antibody against H3K79me2 (Abcam, ab3594, 1:1,000) for 1 hour. Next, the cells were incubated for 1 hour with Alexa Fluor 555-conjugated secondary antibody (ThermoFisher, A-31572, 1:1,000) and DAPI (ThermoFisher, #62249, 1:10,000). Pictures were taken using an Olympus IX83 microscope. Fluorescence quantification was performed with ImageJ Software (NIH Image, National Institutes of Health) using 20 pictures per condition for each independent experiment.

### DMM mouse model of OA

All experiments with mice were approved by the Ethics Committee for Animal Research (KU Leuven, Belgium). Wild-type male C57Bl/6 mice were purchased from Janvier (Le Genest St Isle, France). At 8 weeks of age, post-traumatic osteoarthritis (OA) was induced by the destabilisation of the medial meniscus (DMM). To this end, a mild instability of the knee was obtained by surgical transection of the medial menisco-tibial ligament of the right knee.(32) Sham-surgery served as control. The knees were histologically analysed 12 weeks after surgery.

### Intra-articular IOX2 injections

One week after DMM surgery, mice were intra-articularly injected with IOX2 (0.5 mg/kg) or vehicle (30% PEG400 in PBS) every 10 days for a total of 7 injections. 12 weeks after surgery, the knees were harvested and analysed.

### Histology

Dissected mouse knees were fixed overnight at 4°C in 2% Formaldehyde, decalcified for 3 weeks in 0.5 M EDTA pH 7.5, and embedded in paraffin. All stainings were performed on 5 µm thick sections. Severity of disease was determined by histological scores on hematoxylin-safraninO stained sections throughout the knee (6 sections at 100 µm distance). Cartilage damage and synovitis were assessed based on OARSI guidelines.(59) Osteophytes were scored following an in-house scoring system earlier reported.(9) Pictures were taken using a Visitron Systems microscope (Leica Microsystems GmbH).

### Immunohistochemistry

Immunohistochemistry was performed on 5 µm thick paraffin-embedded EDTA-decalcified knee sections. Heat induced epitope retrieval was performed using a Citrate-EDTA buffer (pH 6.2) for 10 min at 95°C. Sections were treated with 3% H2O2/methanol for 10 min to inactivate endogenous peroxidase, blocked in goat serum for 30 min and incubated overnight at 4°C with primary antibodies against DOT1L (Abcam, ab64077, 6 µg/ml), HIF-1α (Abcam, ab82832, 10 µg/ml) or for 90 min with primary antibody against H3K79me2 (Abcam, ab3594, 1 µg/ml). Rabbit IgG (Santa Cruz, sc-2027) was used as negative control. Avidin-biotin complex amplification (Vectastain ABC kit, Vector Laboratories) was used, except for the immunohistochemical detection of H3K79me2. Peroxidase goat anti-rabbit IgG (Jackson Immunoresearch) was applied for 30 min and peroxidase activity was determined using DAB.

### Micromasses

Primary hACs were cultured in 10 µl droplets (micromasses) in 24-well plates at a density of 300,000 cells/micromass. Culture medium was changed twice per week and consisted of DMEM/F12 (Gibco), 10% FBS (Gibco), 1% (vol/vol) antibiotic/antimycotic (Gibco), 1% L-glutamine (Gibco) and Insulin-Transferrin-Selenium (ITS) (Thermo Fisher). Micromasses were treated with vehicle (DMSO) or DOT1L inhibitor EPZ-5676 (EPZ) (10µM) (Chemietek) under normoxic (21% O_2_) or hypoxic (1% O_2_) conditions for 2 weeks. The micromasses were washed with PBS and fixed with ice-cold methanol for 1h at -20°C. After rinsing with PBS, the micromasses were stained with Alcian Blue (0.1% AB 8GX, Sigma) for 2,5h, washed with water and air dried. Quantification of the staining was performed by dissolving the micromasses with 6 M guanidine (Sigma) for 6h and measuring the absorbance at 595 nm with a spectrophotometer (BioTek Synergy).

### Study Approval

Primary human primary articular chondrocytes (hACs) were isolated from patients undergoing hip replacement surgery for osteoporotic or malignancy-associated fractures with informed consent and ethical approval of The University Hospitals Leuven Ethics Committee and Biobank Committee (Leuven, Belgium) (S56271). According to Belgian Law and UZ Leuven’s biobank policies, the hip joints are considered human biological residual material. Only age and sex of the patients are being shared between the surgeons and the investigators involved in this study. The material is fully anonymised without links to the medical file. All mouse model studies were performed with the approval from the Ethics Committee for Animal Research (P114-2008, P198-2012, P159-2016; KU Leuven, Belgium) (Licence number LA1210189).

### Statistics

Data analysis and graphical presentation were performed with GraphPad Prism version 8. Data are presented as mean and SEM and as individual data points, representing the mean of technical replicates as indicated in the figure legends. Gene expression data and ratio data were log-transformed for statistical analysis. All test performed were two-tailed where applicable. For comparisons against a hypothetical mean in the ChIP experiments, one sample t-test was used For comparisons between two groups, Student’s t-test (unpaired and paired as indicated in figure legends) assuming equal variance was used. For comparisons between more than two groups, one-way ANOVA was used. For dose-response experiments, Linear trend estimation ANOVA was used. Two-way ANOVA (analysis of variances) was used to study interactions and main effects between independent categorical variables. Sidak correction was used for multiple comparisons. Data are reported by F-values and t-values with degrees of freedom and exact *p*-values in the figure legends where applicable. Effect sizes (R^2^ for regression models or differences between means in 2 group comparisons) are reported in Results section. Distribution of the dependent variables was assessed by histogram inspection and by QQ plots. Homogeneity of variance was evaluated by residuals versus fit plot.

## AUTHOR CONTRIBUTIONS

A.D.R., S.M. and R.J.L. planned the study and designed all the in vitro, ex vivo, and in vivo experiments. A.D.R. and A.E. performed in vitro and ex vivo experiments. F.M.F.C. performed the animal experiments. C.C. contributed to experimental design. F.C. cloned and provided the plasmid constructs. R.J.L. and A.D.R. are responsible for all statistical analyses. L.C. and A.S. provided essential materials. A.D.R., S.M. and R.J.L. wrote the manuscript.

## ACKNOWLEDGMENTS

We are grateful to L. Storms for her technical support in this study and for, together with A. Hens, taking care of the animal facility management. We thank I. Jonkers and J. Peeters for making the hypoxia incubator available. We are indebted to the UZ Leuven traumatology and orthopaedics nursing staff for their efforts to provide cartilage samples for ex vivo and in vitro work.

