## Supplement for "Hypoxia induces transcription of DOT1L in articular cartilage to protect against osteoarthritis"

### SUPPLEMENTARY INFORMATION

#### Supplementary figure S1

**A** >DOT1L; human; range -1000bp to 100bp.

TACCTCAGCCCCCGCAGTAGCTGGGATTACAGGCGCCCGCCACCGCGCCCAGCTAATTTTTGTATTTTTA
ATAGAGACAGGGTTTCACCATGTTGGCCAGGCTGATCTCGAACTCCTGACCTCAGGTGATCTTCCCGCCT
CGGCCTCACAAAGTGCTGGGATTACAGGCATGAGCCAGGGCTCCTGGCCGTGAGAGGGGGTTTCTCCATG
TTGGTCAGGCTGGTCTCGAACTCCTGACCTCAGGTGATCTGCCCGCCTTGGCCTCCCAAAGTGCTGGGAT
TACAGGGGTGAATCACCGCGCCTGGCCTTTTTTTTTTTTTTTGTACCCAATAAACAGCATTGTTGTCGAA
TGAATACTGGACTTGGAGCTACAGCAAGTCCTGGACCGTGACTCTTATGGGGGAAATTCGGATTTTTGG
TTTTACTAAGCCGTGTGTGGGGAGGTGTCCGGCGTCCTCCTCCTGGGACGGGATTCGAACCCGCTATCCG
ACGGGCCCGCCCACAGGGTCTCCCCGGGTCCCCGCTTCGGGCCGGCGAGTGGGGGAAGGGGTCGGCCGAG
GGCAACCGAGGACGTGCGTGCGTACGTTCGTGCGTGCGTGGATTCGGGCGGGCGGGCGAGTCCACGGGGC
GGGGCGCCGAGGGGGTGGCGCGCGCGGGTCGGCCCGCTGGGCGGCGGGCACGCGCCGGCGTCTTCGCTCC
GGGCTCCCCTAGCGCGCGGGGCGAGTGGTTCCGCCCGGCCCCCGGCTCATTGTGCTCGCTTCACGCCGGC
CCAAGATGGCGGAGGCGCTGGAGGCCCCGGGCCTGTGACTACAAAGAGGGAGTCGGGGGCCGGGCCGGAC
CGGAGCGCGGCGGCGGCGGCGGCGGCGGCCGAGGCCGAGGCCAGGCCCCCTCCCCTCAGCCTCCCGCCCC
TCCCTCCCGCCCGCCCTCCTCCGCCCACCGGCGGCCCCGCCCCTCCCCCAACCGCCCGCCTAGCATGGTG
CGGCGGCCGCGCGCGCGGACATGGGGGAGAAGCTGGAGCTGAGACTGAAGTCGCCCGTGGGGGCTGAGCC
CGCCGTCTACCCGTGGCCGCTGCCGGTCTACGTGAGTGCCGCCCTCCACCG

**B** >DOT1L; mouse; range -1000bp to 100bp.

TTCTAGCTCCGGATCCTTCTTTTGGGGGCTGGCAAGCGCTAGGATGAGGGGTGGGAGCCATCTCCCCTAG
ATTACATTCACTCCCATGTAGTCAGGAAGAAGAAGCAGCTATCCAGAAGCGGCTGGTGAGCTGGTGACAG
CTCACCTCAAGGTAACTTATTCTCAGAAGGTTTTGGGGGGAGGGAACTTAAGTATTAGAAGGATTGGAAT
GGCCTAAGAAGGAAATAAGTTTGGCTCCTGACTCTGTAAAGTTCTGGGCAGGTCAGAGAATAACATTGCT
TTGAACTCTTAATTGAGAGCCTCCGACCTATAGGGTCCAATTTGACTGATGATTTATTAATTAACTTATT
GCTCACATCTTTAATCCCAGCTCTCAGGAGGCAGAGGCAGGTGGATTTTGGTAAATTCGAGGTCAGACTG
GTCAACAGAATGAGTTCCTGGACAGCCAGGGACACCCTGTCTCAAAAAAAAAAAAAAAAAAAAAAGCATG
TAAATAAACAATAAGCAGATAGGCTAGAAAAGTGAAAAAATAGAAAAGCTAAAGAAGGGAAAAATTGAA
AAATGAAAAAAAAAAGAAAGAAAGAAAAAGCCAGAAAAACAGAAAAAGAAAATTGACGGGAAGGGAAGAG
AAAGAAAAAGAACAGGAAATGGAAAGAAAAGAAAAAAGCAAAGCAAAGCGAAGGGACGCGAAGCAAGCCT
GTGCAACTGGGATTTGAACTTGACCCCGCCTGGTCTCTCTCGCTTTCCTCCGCAGGGTTTCCCCGGGTCC
CCGCTTCCGGCTGGCGGCGACGTCGGAGGGCAACCGAGGACGTGCGTGCGCCGCTGCGTGCGTAAGTGCG
TGCGTCGGTCGGAGGAGGGCGAGTCCAAGGGGGCGTGGCGTGCGAGGGGTGGCGCGCGCGGCGCGGCGGG
CGGAGGCTCTGGGCGCGCACCGCCTCTTGGCTCCCGGCCTCCCCCGCGCGCGCTCGCCGAGTGGTTCCGC
CCGACCCCCCGGCTCATTGTGCCTTCTCCTCACGCCGGCCCAAGATGGCGGCGGCTCTAGACGCCCCGGC
TCTGTGACTCCTACAAAGAGGGGAGCTGGGGCCACACGGGAGCGGCGGCCG

**C** >DOT1L; rat; range -1000bp to 100bp.

ATGTATGTTTATGCACCACGATTGCCTGGTGCATGGAGAGGCCAGAAGAGGGCGTTGGAACCCTGGGAAT
TCAGTCAGAGATGGTTGTGAGCCCGAGGGAGGGCTGGGAATGGAATCTAAGGTTCTCTAAAAGAGCAGCC
TGTAGTCTTAGCTACTGAACCATCTCTCTAACCCATTTGATGATTTTGTTTTTCAGTTTTGTTCTTGCTG
TTAGGAATGGTTTCACTCTGTAGCTCAAGTGGGCTTTGAGTCGTGATCCCTTGCCTCAGCTTCCCAAGAC
CTTGGATTATGGGTCATAGATTCACGGTTTTGTTAGTGTGAAACTGCATAGAAAGTCGGAAAATATCTTC
AAAAAGAATGAAAAGAAGCTAGGCGTGGTGGTGCACACCTTTAATCCCAGCACTAGGGAGGGAGGCAGAG
GCGGGGAATCTTGGTGAATTTGAGGTAAGACTGGTCAACAGAGTAGACGGAGTTCCAGGACAGCCAGGGC
TACCGTGTCTCAAAAAAAAAAAAAAAACCAAATAAGTAAATAAACAATAAATAAGTGGATAAGATAAAAA
AAAGTGAAAAGAGAAACAGAAAAAGGAAAAAAGCCGGAAAAACAGAAAAAGAATAGGGACGAGAAGAGAA
AGAAAAGGAAAAGGAAAGGAAAGAAAAGAAAAAGAAAAGAAAAGAAAAAAAAGCAAAGCAAAGCGAGCCT
GCGCAACTGGGACTTGAACTTGTCCCCACCTGGTCTCTCTCGCTTGCCTCAGCAGGGTTTCCCCGGGTCC
CCGCTTCCGGCTGGCGGTGGCGTCGGAGGGCAACCGAGGACGTGCGTGCGACCCTGCGTGCGAACGTGCG
TGCGTCGGCGGGAGGTGGGCGAGTCCAAGGGGGCGTGGCGTACGAGGGGTGGTGCGCGCGGCGCGGCGGG
CGGAGGCTTTGGGCGCGCACCGCCTCTTGGCTCCCGGCCTCCCCCGCGCGCGCGCGCCCAGTGGTTCAGC
CCGACCCCCCGGCTCATTGTGCCTTCTCCTCACGCCGGCCCAAGATGGCGGCGGCGCTAGACGCCCCGGC
TCTGTGACTCCTACAAAGAGGGGAGCTGGGGCCACACGGGAGCGGTGGCCG

**Supplementary figure S1** Predicted hypoxia response elements (HREs) in human, mouse and rat *DOT1L* gene promoters. (A-B-C) Sequences of the human (A), mouse (B) and rat (C) *DOT1L* gene promoters from 1000 base pairs (bp) upstream of the transcription start site (TSS) to 100 bp downstream of the TSS as determined by the Eukaryotic Promoter Database. HREs with core consensus sequence 5'-(A/G)CGTG-3', potential binding sites for HIF heterodimers, are marked with a yellow box. The bp downstream of the TSS are highlighted in grey.

### Supplementary figure S2

**Multiple sequence alignment by MUSCLE**

```
human -------------TACCTC--------AGCCCCGCAGTAGCTGGGATTACAGGCGCCCGCCACCGCGCCCAG-------CTAATTTTTGTATTTTT
mouse TTCTAGCTCCGGATCCTTCTTTTG---GGGGCTGGCAAGCGCTAGGATGAGGGGTGGGAGCCATCTCCCCTAGATTACATTCACTCCCATGTAGTC
rat   ATGTATGTTTATGCACCACGATTGCCTGGTGCATGGAGAGGCCAGAAGAGGGCGTTGGAA-------CCCTGG---GAATTCAGTCAGAGATGGTT
                     *  *         *  *  * *   **  * *      *             **  *        * * *      *  *

human AATA----GAGACAGGGTTT-----------------------------CACCATGTTGGCCAGGCTG-------ATCTC---------GAACTCC
mouse AGGAAGAAGAAGCAG-----------------------CTATCCAGAAGCGGCTGGTG-----AGCTGGTGA--CAGCTCACCTCAAGGTAACTTA
rat   GTGAGCCCGAGGGAGGGCTGGGAATGGAATCTAAGGTTCTCTAAAAGAGCAGCCTGTAGTCTTAGCTACTGAACCATCTCTC-------TAACCCA
         *    **   **                                  *  *  **       ***        * ***          ***

human TGACCTCAGGTGATCT-------------------------------------------------------------------------------T
mouse T--TCTCAGAAGGTTTTGGGGGGAGGGAACTTAAGTATT-------------AGAAGGATTGGA----ATGGCCTAAGAAGGAAATAAGTTTGGCT
rat   T--TT---GATGATTTTGTT---------TTTCAGTTTTGTTCTTGCTGTTAGGAATGGTTTCACTCTGTAGCTCAAGTGGGCTTTGAGTCGTGAT
      *       *  * * *                                                                               *

human CC--CGCCTCGGCCTCACAAAGTGCTGGGATTACAGG-----------CATGAGCCAGGGCTCCTGGCCGTGAG----------AGGGGGTT----
mouse CC--TGACTCTG-----TAAAGTTCTGGG----CAGGTCAGAGAATAACATTGCTTTGAACTCTTAATTGAGAGCCTCCGACCTATAGGGTC----
rat   CCCTTGCCTCAGCTTC-CCAAGACCTTGGATTATGGGTCATAGATT--CACGGTTTG-------TTAGTGTGAAACT---GCATAGAAAGTCGGAA
      **   * *** *       ***  ** **      **           **       *      *    * **           *    **

human ----TCTCCATGT-----------------------TGGTCAGGCTGGTCTCGAACTCCTGACCT-----------------CAGGTGATCTGCCC
mouse --------CAATTTGACTGATGATTTATTAATTAACTTATTGCTCACATCTTTAATCCCAGCTCT----CAGGAGGCAGAGGCAGGTGG-------
rat   AATATCTTCAAAAAGAATGAAAAGAAGCTAGGCG--TGGTGGTGCACACCTTTAATCCCAGCACTAGGGAGGGAGGCAGAGGCGGG-GA-------
              **                          *  *    *    **  **  ** *  **                 * ** *

human GCCTTGGCCTCCCAAAGTGCTGGGATTACAGGGGT--------------GAATCACCG---CGCCTGGCCTTTTTTTTTTTT--------------
mouse ATTTTGG-----TAAATT--CGAGGTCAGACTGGTCAACAGAAT-----GAGTTCCTGGACAGCCAGGGACACCCTGTCTCAAAAAAAAAAAAAAA
rat   ATCTTGG-----TGAATT--TGAGGTAAGACTGGTCAACAGAGTAGACGGAGTTCCAGGACAGCCAGGGCTACCGTGTCTCAAAAAAAAAAAAAAA
         ****       ** *   * * * * *  ***              ** *  * *    *** **       * * *

human --------TTTGTACCCAATAAACAGCA----------------TTGTTGTCGAATGAATA--CTGGACTTGGAGCTACAGCAAGTCCTGGACCGT
mouse AAAAAAAGCATGTA---AATAAACAATAAGCAGATAGGCTAGAAAAGTGAAAAAATAGAAAAGCTAAAGAAGGGGAAAAATTGAAAAATGAAAAAA
rat   ACCAAA--TAAGTA---AATAAACAATA---------------------AATAAGTGGATAAGATAAA-------AAAAAGTGAA-----AAGAGA
         ***   ********  *       *                         * *  * *   *  *         * *   *       *

human GACTCTTATGGGGGGAAATTCGGATTTTTGGTTTTACTAAGCCGTGTGTGGGGAGGTG--------------------------------------
mouse AAAAGAAAGAAAGAAAAAGCCAGAAAAACAG----AAAAAGAAAATTGACGGGAAGGGAAGAGAAAGAAAAAGAACAGGAAATGGAAAGAAAAGAA
rat   AACAGAAAAAGGAAAAAAGCCGGAAAAACAG----AAAAAGAATAGGGACGAGAAGAGAAAGAAAAGGAAAAGGAAAGGAAA--GAAAAGAAAAAG
       *     *       ***  * **      *    *   ***     *  * ** * *

human -----------------------TCCGGCGTCCTCCTCCTGGGACGGGATTCGAACCCGCTATCCGACGGGCC-----------CGCCCACAGGGT
mouse AAAAGCAAAGCAAAGCGAAGGGACGCGAAGCAAGCCTGTGCAACTGGGATTTGAACTTGAC-CCCGCCTGGTCTCTCTCGCTTTCCTCCGCAGGGT
rat   AAAAGAAAAG-AAAAAAAAGCAAAGCAAAGCGAGCCTGCGCAACTGGGACTTGAACTTGTC-CCCACCTGGTCTCTCTCGCTTGCCTCAGCAGGGT
                               *   *    ***        **** * ****  *    **  * ** *           *  *  ******

human CTCCCCGGGTCCCCGCTTCGGGCCGGCGAGTGGGGAAGGGGTCGGCCGAGGGCAACCGAGGACGTGCGTGCG----TACGTTCGTGCGTGCGTGG
mouse TTCCCCGGGTCCCCGCTTCCGGCTGGC-----------GGCGACGTCGGAGGGCAACCGAGGACGTGCGTGCGCCGCTGCGTGCGTAAGTGCGTGC
rat   TTCCCCGGGTCCCCGCTTCCGGCTGGC-----------GGTGGCGTCGGAGGGCAACCGAGGACGTGCGTGCGACCCTGCGTGCGAACGTGCGTGC
       ****************** *** ***           ** * ** * *************************    * *** **   *******

human ATTCG-GGCGGGCGGGCGAGTCCAC-GGGGCGGGGCGCCGAGGGGGTGGCGCGCGCGGGTCGGCCCGCTG-GGCGGCGGGCACGCGCCGGCGTCTT
mouse GTCGGTCGGAGGAGGGCGAGTCCAAGGGGGCGTGGCGTGCGAGGGGTGGCGCGCGCGGCGCGGCGGGCGGAGGCTCTGGGCGCGCACC-GCCTCTT
rat   GTCGGCGGGAGGTGGGCGAGTCCAAGGGGGCGTGGCGTACGAGGGGTGGTGCGCGCGGCGCGGCGGGCGGAGGCTTTGGGCGCGCACC-GCCTCTT
       *  *  *  ** ***********  ****** ****     ******* ********  ****  ** * ***   **** *** ** ** ****

human CGCTCCGGGCTCCCCTAGCGCGCG--GGGCGAGTGGTTCCGCCCGG-CCCCCGGCTCATTGTGC--TCGCTTCACGCCGGCCCAAGATGGCGGAGG
mouse GGCTCCCGGCCTCCCCCGCGCGCGCTCGCCGAGTGGTTCCGCCCGACCCCCCGGCTCATTGTGCCTTCTCCTCACGCCGGCCCAAGATGGCGGCGG
rat   GGCTCCCGGCCTCCCCCGCGCGCGCGCGCCCAGTGGTTCAGCCCGACCCCCCGGCTCATTGTGCCTTCTCCTCACGCCGGCCCAAGATGGCGGCGG
       ***** ***  ***  *******   * * ******** *****  *****************  ** * ********************** **

human CGCTGGAGGCCCCGGGCCTGTGA---CTACAAAGAGGGAGTCGGGGGCCGGGCCGGACCGGAGCGCGGCGGCGGCGGCGGCGGCGGCCGAGGCCGA
mouse CTCTAGACGCCCCGGCTCTGTGACTCCTACAAAGAGGGGAGCTGGGGCCACACGGGA---------------------------------------
rat   CGCTAGACGCCCCGGCTCTGTGACTCCTACAAAGAGGGGAGCTGGGGCCACACGGGA---------------------------------------
      * ** ** *******  ******   ************   * ******   * ***

human GGCCAGGCCCCCTCCCCTCAGCCTCCCGCCCCTCCCTCCCGCCCGCCCTCCTCCGCCCACCGGCGGCCCCGCCCCTCCCCCAACCGCCCGCCTAGC
mouse ------------------------------------------------------------------------------------------------
rat   ------------------------------------------------------------------------------------------------


human ATGGTGCGGCGGCCGCGCGCGCGGACATGGGGGAGAAGCTGGAGCTGAGACTGAAGTCGCCCGTGGGGGCTGAGCCCGCCGTCTACCCGTGGCCGC
mouse -----GCGGCGGCCG---------------------------------------------------------------------------------
rat   -----GCGGTGGCCG---------------------------------------------------------------------------------
           **** *****

human TGCCGGTCTACGTGAGTGCCGCCCTCCACCG
mouse -------------------------------
rat   -------------------------------
```

**Supplementary figure S2** Alignment of the human, mouse and rat *DOT1L* gene promoters. Alignment of the human, mouse and rat sequences of the *DOT1L* gene promoters from 1000 base pairs (bp) upstream of the transcription start site (TSS) to 100 bp downstream of the TSS as determined by the Eukaryotic Promoter Database. This alignment was performed using MUSCLE. Hypoxia response elements (HREs) are marked with a yellow box.

**Supplementary figure S3**

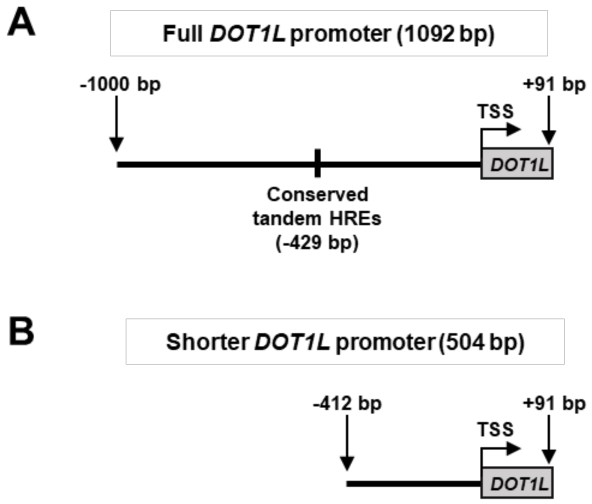


**Supplementary figure S3** Promoter constructs for the luciferase reporter assay. (A) The full human *DOT1L* promoter from 1000 base pairs (bp) upstream to 91 bp downstream relative to the transcription start site (TSS) as defined by the Eukaryotic Promoter Database was cloned into the pGL3 basic luciferase reporter vector. This 1092 bp fragment contains the conserved overlapping tandem Hypoxia response elements (HREs) located at -429 bp relative to the TSS. (B) The shorter *DOT1L* promoter from 412 bp upstream to 91 bp downstream relative to the TSS was cloned into the pGL3 basic luciferase reporter vector. This 504 bp fragment does not contain the conserved overlapping tandem HREs.

**Supplementary figure S4**

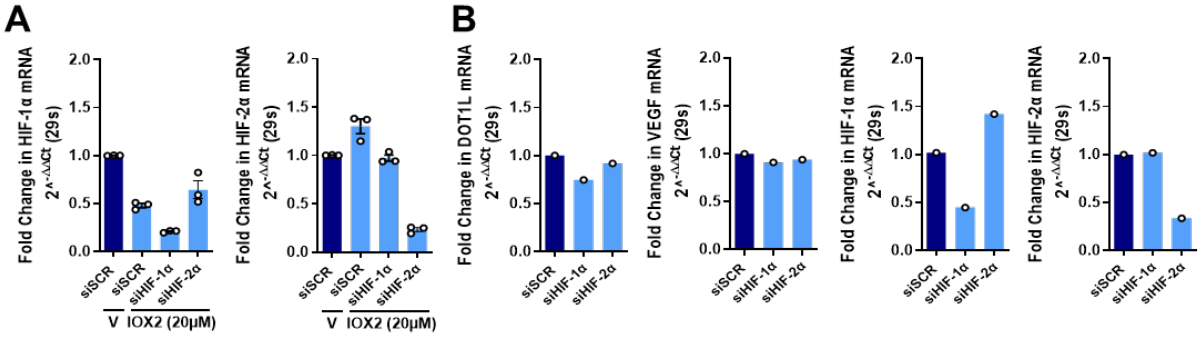


**Supplementary figure S4** Efficiency of siRNA-mediated gene silencing in C28/I2 cells. (A) Real-time PCR analysis of *HIF-1α* and *HIF-2α* in human articular chondrocyte C28/I2 cells after treatment with hypoxia mimetic IOX2 (20 μM) and siRNA-mediated silencing of *HIF-1α* (siHIF-1α), *HIF-2α* (siHIF-2α) or scrambled control (siSCR) (n=3, $t_8$= 7.013, 7.786, 2.639 – $p$=0.0003, =0.0002, =0.0867 for *HIF-1α*; $t_8$= 3.208, 3.617, 21.57 – $p$=0.0369, =0.203, <0.0001 for *HIF-2α*; V-siSCR vs IOX2-siSCR, IOX2-siSCR vs siHIF-1α, siHIF-1α vs siHIF-2α, Sidak-corrected in one-way ANOVA with $F_{3,8}$=77.26 $p$<0.0001 for *HIF-1α* $F_{3,8}$=191.4 $p$<0.0001 for *HIF-2α*). (B) Real-time PCR analysis of *DOT1L*, *VEGF*, *HIF-1α* and *HIF-2α* in C28/I2 cells after siRNA-mediated silencing of *HIF-1α*, *HIF-2α* or scrambled control (siSCR) (n=1).

**Supplementary figure S5**

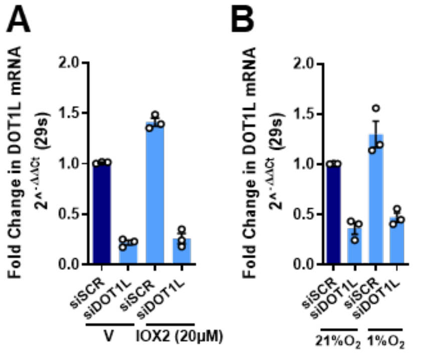


**Supplementary figure S5** Efficiency of siRNA-mediated gene silencing in hACs. (A-B) Real-time PCR analysis of *DOT1L* in primary human articular chondrocytes (hACs) after treatment with hypoxia mimetic IOX2 (20 μM) or vehicle (V) (A) (n=3, $t_8$=9.678 $p$<0.0001, $t_8$=10.86 $p$<0.0001 siSCR vs siDOT1L with V and IOX2 for *DOT1L* Sidak-corrected in two-way ANOVA with $F_{1,8}$=4.450 $p$=0.0679 for treatment status, and $F_{1,8}$=210.9 $p$<0.0001 for silencing status) or culturing in hypoxic conditions (1% $O_2$) (B) and siRNA-mediated silencing of *DOT1L* or scrambled control (siSCR) (n=3, $t_8$=6.605 $p$=0.0003, $t_8$=6.360 $p$=0.0004 siSCR vs siDOT1L in normoxia and hypoxia for *DOT1L* Sidak-corrected in two-way ANOVA with $F_{1,8}$=5.552 $p$=0.0462 for oxygen status, and $F_{1,8}$=84.05 $p$<0.0001 for silencing status).

**Supplementary figure S6**

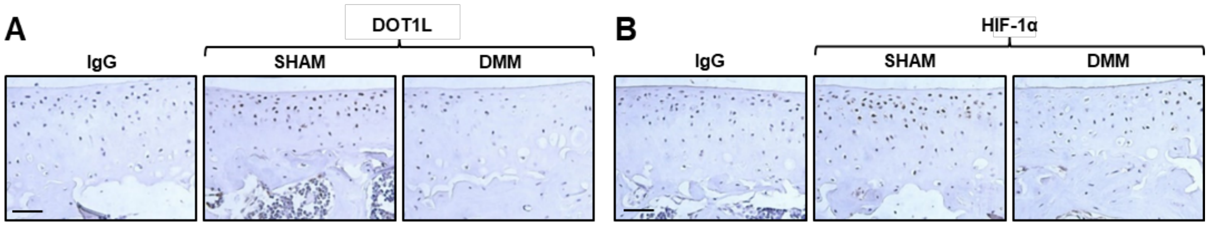


**Supplementary figure S6** DOT1L and HIF-1α protein levels in osteoarthritis murine articular cartilage. (A-B) Immunohistochemical detection of DOT1L (A) and HIF-1α (B) in the articular cartilage of wild-type mice with OA triggered by destabilisation of the medial meniscus (DMM) surgery compared to sham operated mice. The images are representative of three different animals. Scale bar, 50 μm.

**Supplementary table S1**

**Human primers used for qPCR**

| Primer name | Sequence |
|---|---|
| **hACAN_fw** | ATCCGAGACACCAACGAGAC |
| **hACAN_rv** | CACTCATTGGCTGCTTCCTG |
| **hCOL2A1_fw** | TGGCAGAGATGGAGAACCTG |
| **hCOL2A1_rv** | CATCAAATCCTCCAGCCATC |
| **hDOT1L_fw** | GGATCTCAAGCTCGCTATGG |
| **hDOT1L_rv** | GTCGATGGCACGGTTGTACT |
| **hHIF1A_fw** | TCATCCATGTGACCATGAGG |
| **hHIF1A_rv** | TTCCTCGGCTAGTTAGGGTACA |
| **hHIF2A _fw** | CTGCGACCATGAGGAGATTC |
| **hHIF2A _rv** | GTACGGCCTCTGTTGGTGAC |
| **hS29_fw** | GGGTCACCAGCAGCTGTACT |
| **hS29_rv** | AAACACTGGCGGCACATATT |
| **hTCF1_fw** | CCCCCAACTCTCTCTCTACGA |
| **hTCF1_rv** | TGCCTGAGGTCAGGGAGTAG |
| **hVEGF_fw** | TGCAGATTATGCGGATCAAACC |
| **hVEGF_rv** | TGCATTCACATTTGTTGTGCTGTAG |

**Supplementary table S2**

#### Human primers used for ChIP-qPCR

| Primer name | Sequence |
|---|---|
| **hDOT1Lprom_fw** | CCAATAAACAGCATTGTTGTCG |
| **hDOT1Lprom_rv** | CCCACACACGGCTTAGTAAAA |
| **hVEGFprom_fw** | TCACTTTCCTGCTCCCTCCT |
| **hVEGFprom_rv** | GCAATGAAGGGGAAGCTCGA |

**Supplementary table S3**

**Primers used for the PCR amplification for luciferase assay**

| Construct | Primer | Sequence |
|---|---|---|
| **Full DOT1L promoter (1092 bp)** | Forward | GGTACCTACCTCAGCCCCCGCAGTA |
| | Reverse | ACGCTCGAGGCGGCACTCACGTAGACC |
| **Shorter DOT1L promoter (504 bp)** | Forward | GGTACCCGTGCGTGCGTGGATTCG |
| | Reverse | ACGCTCGAGGCGGCACTCACGTAGACC |
